# Superresolution architecture of pluripotency guarding adhesions

**DOI:** 10.1101/402305

**Authors:** Aki Stubb, Camilo Guzmán, Elisa Närvä, Jesse Aaron, Teng-Leong Chew, Markku Saari, Mitro Miihkinen, Guillaume Jacquemet, Johanna Ivaska

**Affiliations:** Turku Centre for Biotechnology, University of Turku and Åbo Akademi University, Turku, Finland; Advanced Imaging Center, HHMI Janelia Research Campus, Ashburn, Virginia 20147, USA; Department of Biochemistry, University of Turku, Turku, Finland

## Abstract

Human pluripotent stem cells (hPSC) can generate almost all adult cell lineages. While it is clear that key transcriptional programmes are important elements for maintaining pluripotency, the equally essential requirement for cell adhesion to specific extracellular matrix components remains poorly defined. Our recent observation that hPSC colonies form unusually large “cornerstone” focal adhesions (FA), distinct from parental somatic cells, that are lost following differentiation, emphasises the potential of these atypical FA as gatekeepers of pluripotency. Here, using nanopatterns, we further demonstrate that physical restriction of adhesion size, in hPSC colonies, is sufficient to trigger differentiation. Using superresolution two-colour interferometric photo-activated localization microscopy (iPALM), we examined the three-dimensional architecture of these cornerstone adhesions and report vertical lamination of FA proteins with three main structural peculiarities: 1) integrin β5 and talin are present at high density, at the edges of cornerstone FA, adjacent to a vertical kank-rich protein wall. 2) Vinculin localises higher than expected with respect to the substrata and displays a head-above-tail orientation, and 3) surprisingly, actin and α-actinin are present in two discrete layers, a previously undescribed localisation for these proteins. Finally, we report that depletion of kanks diminishes FA patterning, and actin organisation within the colony, indicating a key role for kanks in hPSC colony architecture.

## Introduction

Human pluripotent stem cells (hPSC) have the capacity to self-replicate and differentiate into virtually any cell type of the human body. This combined with the seminal finding that hPSC can be generated by artificial reprogramming of somatic cells (Takahashi et al., 2007; Yu et al., 2007), underscores the immense therapeutic potential of these cells. The delineation of transcriptional programs that control pluripotency as well as the development of protocols to induce specific differentiation programs are at the forefront of research to harness the power of hPSC (Boyer et al., 2005). Towards this end, a key consideration for the safe clinical use of hPSC, is the requirement for animal serum- and feeder-layer-free culture conditions that support pluripotency. Important discoveries, made in the past decade, demonstrate that, in vitro, hPSC adhesion to specific extracellular matrix (ECM) molecules is fundamental for maintaining cells in a pluripotent state (Braam et al., 2008; Rodin et al., 2014; Baxter et al., 2009), thus highlighting a fundamental role for the ECM, as well as ECM receptors, for stemness.

Cells interact with the surrounding ECM via transmembrane adhesion receptors, such as integrins, which provide a physical link between the ECM and the actin cytoskeleton (Askari et al., 2009). Integrin engagement leads to the assembly of large signalling platforms termed focal adhesions (FA), which modulate most cellular functions (Legate et al.,2009) FA are highly dynamic and complex structures composed of hundreds of proteins (Horton et al., 2015, 2016) that tune cellular responses, but their overall architecture can be studied using a handful of key adaptors. Indeed, advances in superresolution light microscopy have enabled the determination of the precise nanoscale architecture of FA in cancer cells. Previous studies have demonstrated highly organized vertical stratification of FA (Kanchanawong et al., 2010; Liu et al., 2015; Case et al., 2015) using a technique that combines interferometry and photoactivated localization microscopy (iPALM) to provide sub 20-nm, three-dimensional (3D) isotropic localization of tagged proteins (Shtengel et al., 2009). FA proteins localize to three general functional layers: a membrane proximal integrin signalling layer (composed of integrins, paxillin and focal adhesion kinase), an intermediate force transduction layer (composed of talin and vinculin) and an actin regulatory layer (composed of both actin and actin regulatory elements) (Kanchanawong et al., 2010; Liu et al., 2015; Case et al., 2015). Interestingly, vinculin a binding partner for many FA proteins and a key stabilizer of talin unfolding has been shown to display a wide distribution across all three layers with increasing upward positioning in mature FA (Case et al., 2015).

Even though integrins are key requirements for maintaining pluripotency in vitro (Rodin et al., 2014; Braam et al., 2008; Meng et al., 2010), hPSC FA composition and organisation remain poorly defined. Previously, we described that the hallmark sharp hPSC colony edge morphology is regulated by unusually large FA, termed cornerstone adhesions, connected by an actin “fence” defined by strong contractile ventral stress fibres (Narva et al., 2017). Furthermore, these colony edge adhesions exert strong traction forces on the underlying ECM, which regulates colony morphology and the orientation of cell division (Narva et al., 2017).

Here, we have used nano-grated patterns to investigate the link between FA morphology and pluripotency and a combination of high- and superresolution microscopy techniques including, iPALM, structured illumination microscopy (SIM) and Airyscan to determine the 3D architecture of cornerstone FA present at the periphery of hPSC colonies. We find that mere physical restriction of FA size, and orientation, in hPSC colonies is sufficient to compromise SOX2 and SSEA-1 expression, indicating a key role for cornerstone FA architecture as “guarding” pluripotency. In addition, we find that the nanoscale localization of several of the key FA components: integrins αV and β5, paxillin, vinculin, talin, actin and α-actinin, proteins previously mapped in classical FA (Kanchanawong et al., 2010), is markedly distinct in hPSC FA. Furthermore, we included novel FA scaffold partners kankl and kank2 in our imaging, and describe a key role for kanks in regulating hPSC adhesion and maintaining FA architecture in pluripotent cells.

## Results

### hPSC cornerstone FA architecture is essential for maintaining pluripotency

hPSC colonies plated on vitronectin (VTN), an ECM ligand which supports pluripotency (Braam et al., 2008), assemble into tightly packed colonies with well-defined edges (Chen et al., 2011). Structurally, these colonies are encircled by unusually large paxillin-positive “cornerstone” FA that are themselves connected by prominent actin bundles termed “actin fence” (Fig. 1a and Supplementary Fig. 1a, (Närvä et al., 2017)). Live-cell imaging of endogenously tagged paxillin revealed that hPSC cornerstone FA are remarkably stable compared to FA formed at the centre of hPSC colonies (Supplementary Fig. 1b, video 1). Structured illumination microscopy imaging further revealed that cornerstone FA are composed of multiple linear units clustered together (Supplementary Fig. 1c).

**Fig. 1.**
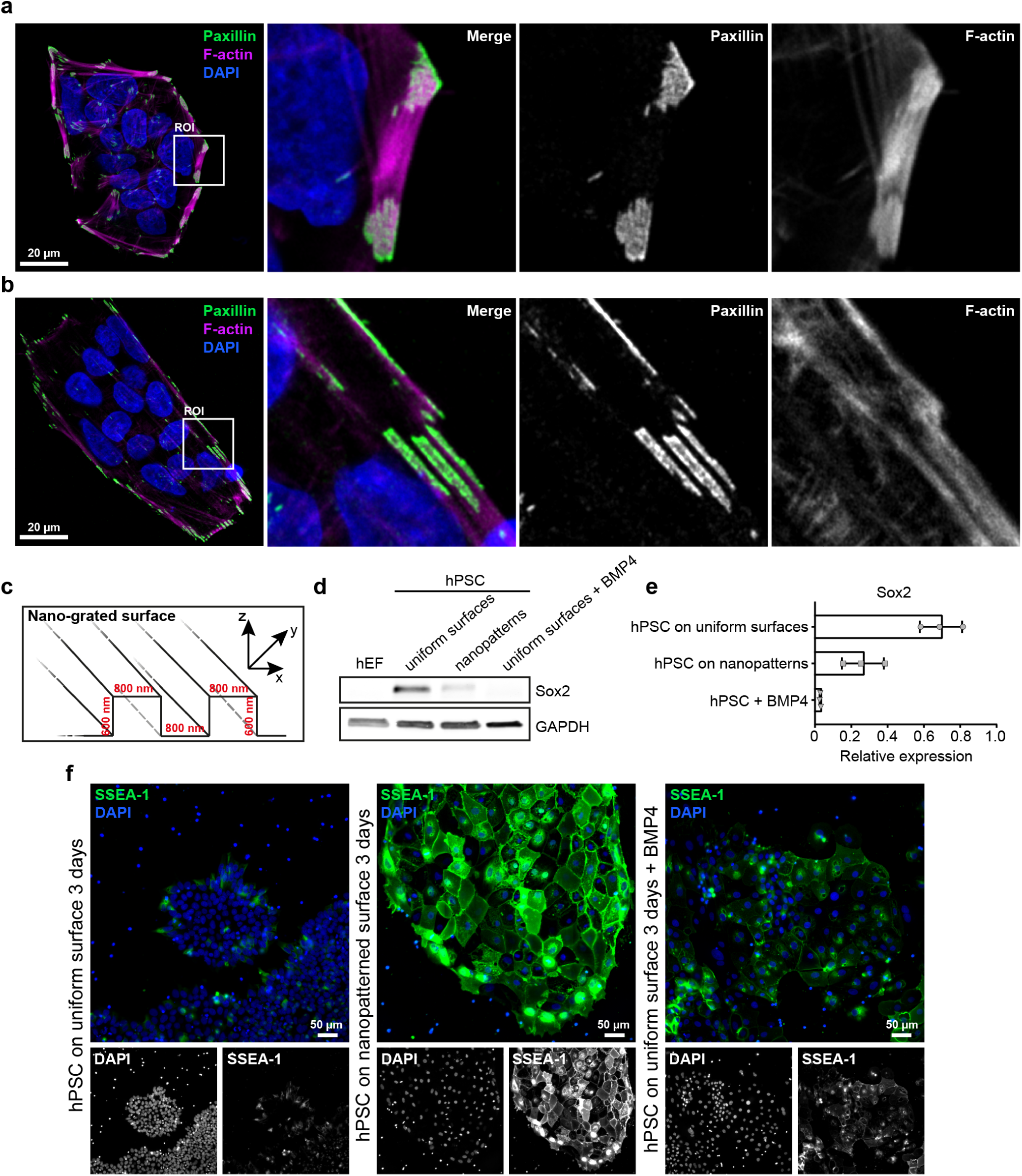
Constraining cornerstone adhesions compromises pluripotency. (a,b) hPSC plated for 24 h on a VTN-coated uniform surface (a) or on a VTN-coated nanograted surface (nanopatterns) (b) in normal (E8 medium) culture conditions, were stained for F-actin, paxillin and DAPI. Images were acquired using an Airyscan confocal microscope. Scale bars 20 μm. A region of interest (ROI) is magnified in each image. (n=3) (c): Cartoon highlighting the topography of the nano-grated surface. (d,e) Representative western blot (d) and quantification (e) of Sox2 levels in human embryonic fibroblasts(hEF) and in hPSC plated for three days either on VTN-coated uniform surfaces, in the presence or absence of BMP4, or on VTN-coated nanopatterns in basal E6 medium (n = 3). (f) hPSC plated for three days either on VTN-coated uniform surfaces, in the presence of basal E6 medium and/or BMP4, or on VTN-coated nanopatterns, in the presence of basal E6 medium, were stained for SSEA-1 and DAPI. Images were acquired on a spinning disk confocal microscope using a 20x objective (n = 3). Scale bars 50 μm.

The unique properties of cornerstone FA (size, orientation and stability) led us to speculate that these structures are key regulators of colony morphology and could contribute to pluripotency maintenance. To constrain the orientation and the maximal size of FA, hPSC colonies were plated on VTN-coated nano-grated patterns of alternating 800 nm wide matrix ridges and 600 nm deep grooves (Fig. 1b and 1c). These nanopatterns were sufficient to constrict adhesion size, reducing the FA characteristic area from 4.2 μm^2^ in cells plated on uniform surfaces, to 1.8 μm^2^ in cells on nanopatterns. Remarkably, on nanopatterns (Fig. 1b and 1c), the characteristic rounded morphology, actin fence and cornerstone FA of hPSC colonies were also distorted (Fig. 1a and 1b). To allow for spontaneous differentiation, hPSC were maintained in E6 medium lacking pluripotency supporting growth factors bFGF and TGFβ. Under these conditions, three-days of exposure to VTN-coated nanopatterns was sufficient to decrease the level of the pluripotency factor Sox2 and, conversely, increase the level of the differentiation marker SSEA-1 (Fig. 1d, 1e and 1f). In contrast, cells maintained on uniform VTN-coated surfaces remained Sox2 positive and SSEA-1 low (Fig. 1d, 1e and 1f). Strikingly, the efficacy of the nanopattern-induced differentiation was comparable to BMP-4 treatment of hPSC colonies on uniform surfaces, which was used as a positive control for differentiation (Fig. 1d, 1e and 1f). These data clearly demonstrate that physical constriction of FA size and orientation in hPSC colonies is alone sufficient to compromise pluripotency and induce differentiation. Importantly, these data indicate a critical role for cornerstone FA architecture in guarding pluripotency, and prompted us to investigate their nanoscale architecture in more detail.

### iPALM imaging of hPSC cornerstone focal adhesions

To deconstruct the ultrastructure of hPSC cornerstone FA, we used iPALM, a strategy that provides sub 20-nm 3D localization of molecules (Shtengel et al., 2009). Previous studies employing iPALM for analysis of osteosarcoma cell (U2OS) adhesions, uncovered FA architecture as a well-organized hierarchy of at least three distinct, vertically separated, functional layers (the integrin signalling layer, the force transduction layer and the actin regulatory layer) (Kanchanawong et al., 2010). To assess the organisation of hPSC cornerstone FA, representative proteins were chosen from each of these layers for imaging, including VTN-binding integrins αV and β5, and paxillin (integrin signalling layer), talin-1 and vinculin (core components of the force transduction layer), and α-actinin-1 and actin (actin regulatory layer) (Kanchanawong et al., 2010). In addition, both kank1 and kank2, two recently described FA-associated proteins, were included due to their unique localisation at the outer-rim of FA (Sun et al., 2016; Bouchet et al., 2016). Of note, the long-isoform of kank1 (Bouchet et al., 2016) is highly expressed in hPSC (Supplementary Fig. 2), and was therefore selected over other isoforms for imaging. Importantly, we could confirm both the endogenous expression of the selected panel of proteins in hPSC (Supplementary Fig. 2) and localisation to cornerstone FA and/or the actin cytoskeleton (Supplementary Fig. 3).

**Fig. 2.**
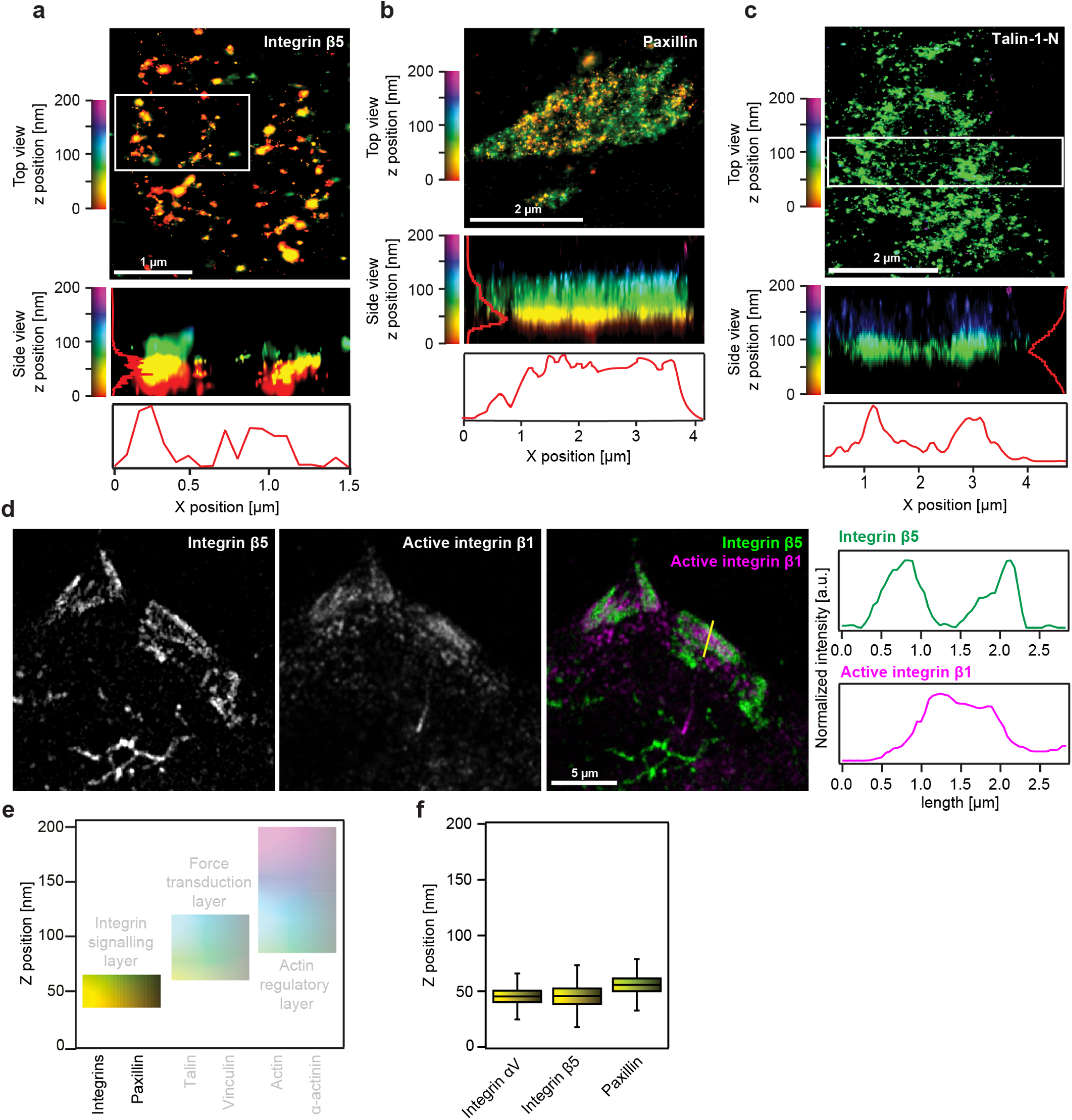
Lateral and vertical segregation of proteins within cornerstone FA. (a-c) Interferometric photo-activated localization microscopy (iPALM) images of Eos-tagged integrin β5 (a), paxillin (b) and talin-1-N (N-terminally tagged talin-1) (c) in cornerstone FA. Both top-view (xy) and side-view (xz) images are displayed. White boxes indicate the area used to generate the side-view images (in the absence of a white box, the entire image was used). For all images, individual localisations are colour-coded as a function of their z-positioning (distance from the coverslip). (d) hPSC plated for 24 h on VTN were stained for endogenous integrin β5 and active integrin β1 (mAb 12G10) and imaged using an Airyscan confocal microscope. The yellow line highlights the area used to measure the intensity profiles displayed on the side. (e) Schematic representation of the different FA layers previously identified using iPALM in U2OS cells (Kanchanawong et al., 2010), highlighting the reported z range for the integrin signalling layer. (f) iPALM analysis of the z position (distance from the coverslip, Z_centre_) of some of the integrin signalling layer components, Eos-tagged integrins αV and β5, and Eos-tagged paxillin, in hPSC cornerstone adhesions. Boxes display the median, plus the 1st and 3rd quartiles (IQR: 25th-75th percentile). Whiskers correspond to the median ± 1.5 x IQR.

**Fig. 3.**
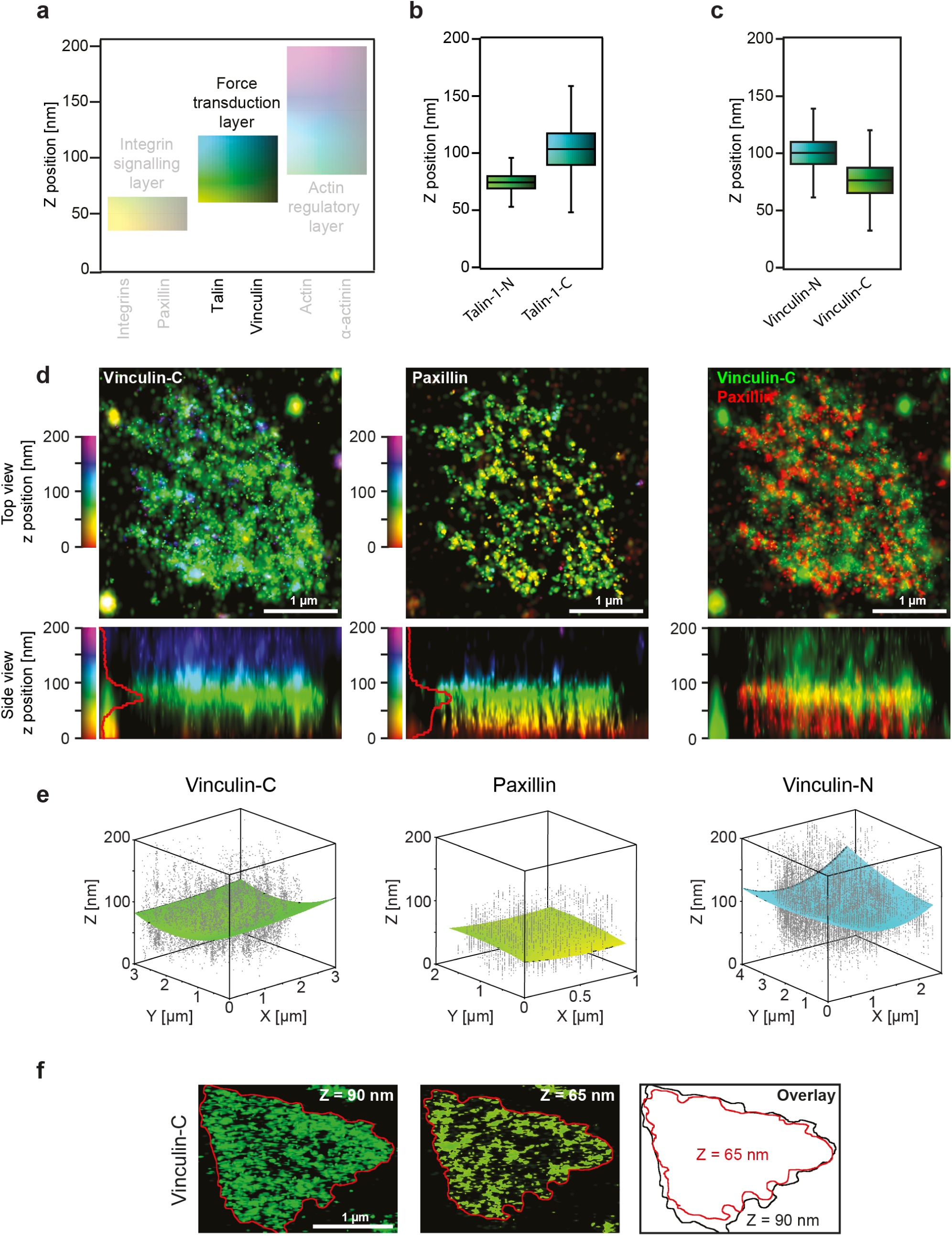
Vinculin distribution and orientation within cornerstone adhesions. (a) Schematic representation of the different FA layers previously identified using iPALM in U2OS cells as in figure 2e, highlighting the reported z range for the force transduction layer containing vinculin and talin. (b) iPALM analysis of the Z_centre_ for N-terminally (Talin-1-N) and C-terminally (Talin-1-C) tagged (Eos) talin-1 in hPSC cornerstone adhesions. Results are displayed as boxplots as in figure 2f. (c) iPALM analysis of the Z_centre_ of N-terminally (Vinculin-N) and C-terminally (Vinculin-C) tagged (Eos) vinculin in hPSC cornerstone adhesions. Results are displayed as boxplots as in figure 2f. (d) Two-colour iPALM images of Eos-taggedvinculin (Vinculin-C) and endogenous paxillin in acornerstone FA. Where localisation ofvinculin and paxillin are displayed separately, top-view and side-view images are colour-coded as a function of the z-position of the indicated protein. Where fluorescence channels are merged (paxillin, red; vinculin, green), the z position is only displayed in the side view and the colours represent the fluorescence signal for each protein. Scale bar 1 μm. (e) 3D scatter plots displaying the individual iPALM localizations (grey dots) of endogenous paxillin and Eos-tagged Vinculin-N and Vinculin-C within a single cornerstone adhesion. Surface plots present the fit of those localisations using a two-dimensional polynomial equation. Note that the paxillin localisations are homogeneously flat while localisations of both vinculin constructs form a solid paraboloid. (f) iPALM xy images of Eos-vinculin-C at selected z-layers in an individual cornerstone FA. Scale bar 1 μm.

Imaging of endogenous molecules by iPALM is however challenging, given the requirement for photo-switchable labels as well as high density labelling (Shtengel et al., 2009). Therefore, and as previously described by others in different cell types (Kanchanawong et al., 2010; Case et al., 2015; Liu et al., 2015), for the purpose of gaining superresolution detail into hPSC FA, proteins of interest, tagged with the photoactivatable fluorescent protein (PA-FP) Eos, were transiently expressed in hPSC. In addition, to image kank1 and kank2, Eos-tagged constructs were generated (see methods for details). We then validated that all Eos-tagged proteins exhibited similar localisation to the endogenous proteins and that the transient expression of these constructs did not affect colony morphology or levels of the pluripotency marker Oct4 (Supplementary Fig. 4).

**Fig. 4.**
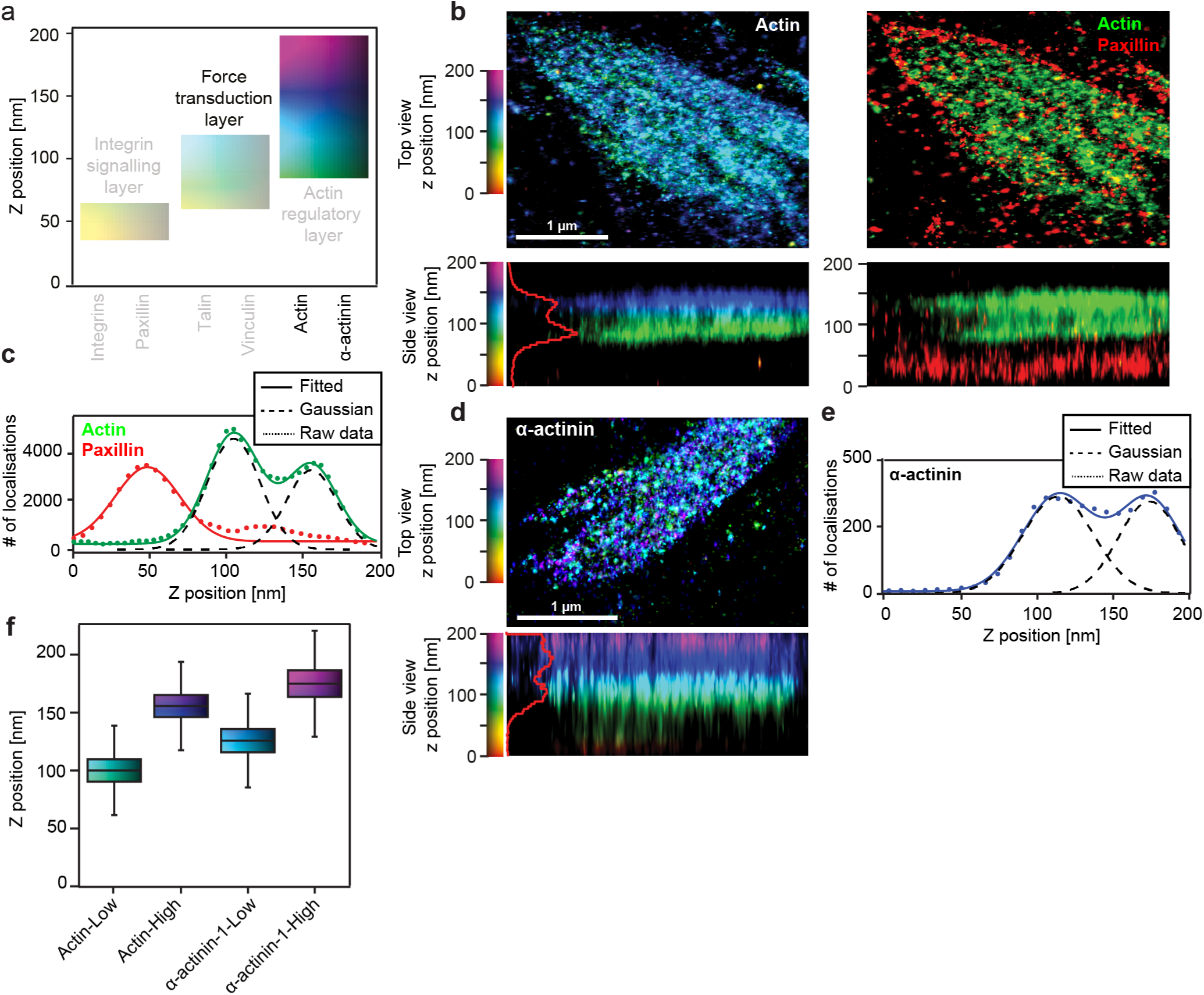
Actin and α-actinin-1 lateral distribution. (a) Schematic representation of the different FA layers previously identified using iPALM in U2OS cells as in figure 2e, highlighting the reported z range for the actin regulatory layer containing actin and α-actinin-1. (b) Two-colour iPALM images of Eos-tagged actin and endogenous paxillin in a cornerstone FA. Where localisation of actin is displayed separately, top-view and side-view images are colour-coded as a function of the z-position of the actin molecules. Where fluorescence channels are merged (paxillin, red; actin, green), the z position is only displayed in the side view and the colours represent the fluorescence signal for each protein. Scale bar 1 μm. (c) Z density profile of paxillin (red) and actin (green) displaying the number of localisations as a function of the z position in an individual cornerstone adhesion. Dotted lines correspond to the experimental data, while solid lines correspond to the fitted data obtained using either a single Gaussian distribution (paxillin) or a sum of two Gaussian distributions (actin). Dashed black lines highlight these two Gaussian distributions. (d) iPALM image of Eos-tagged α-actinin-1 in an individual hPSC cornerstone FA. Scale bar 1 μm. (e) Z-density profile of α-actinin-1 (purple) showing the number of localisations as a function of the z position. Dotted line corresponds to the experimental data while the solid line corresponds to the fitted data obtained using a sum of two Gaussian distributions (dashed black lines). (f) iPALM analysis of the Z_centre_ of actin and α-actinin-1 in hPSC cornerstone adhesions. “Low” and “High” denote separate peaks in the distribution of the same protein. Results are displayed as boxplots as in figure 2f.

### Talin-1 and integrin β5 accumulate at the periphery of large cornerstone adhesions

Using iPALM, we first determined the lateral distribution of proteins localising to the various FA functional layers. All proteins, with the exception of β5 integrin and talin-1, displayed homogeneous x-y distribution and density within cornerstone FA. On the other hand, β5 integrin and talin-1 revealed an obvious ring-like distribution, with higher protein density at the edges of the cornerstone adhesions (Fig. 2a-c). The integrin *α* subunit partner for β5, αV, was, however, homogenously distributed, likely reflecting the known interaction of αV with multiple other β-integrin subunits (Humphries et al., 2006).

The accumulation of Eos-tagged β5 integrin and talin to the rim of the large cornerstone FA was recapitulated with staining of the endogenous proteins imaged on an Airyscan confocal microscope (Supplementary Fig. 5a and 5b). In this case, αβ5 integrin (detected either with an antibody recognizing the β5 subunit or the αVβ5 heterodimer) was strongly localised in FA edges (Fig. 2d and Supplementary Fig.5c), while active β1 integrin and paxillin were distributed uniformly throughout the interior of the adhesion (Fig. 2d and Supplementary Fig.5d). Thus, in hPSC cornerstone FA, the integrin signalling layer appears to be horizontally segregated into sub-regions of different VTN-binding integrins.

### Talin is fully extended throughout cornerstone focal adhesions

We next used iPALM to determine the vertical z positioning of the chosen adhesion proteins (distance measured from the coverslip) (Fig. 2e). We found that the components of the integrin signalling layer (integrins β5 and αV, and paxillin) have a similar vertical distribution in hPSC cornerstone FA to those reported for U2OS FA with z-positions indicating a close relationship with the cell membrane (Fig. 2f). However, talin-1 and vinculin, components of the force transduction layer, exhibited a higher z-position than previously observed (Kanchanawong et al., 2010; Case et al., 2015; Liu et al., 2015) (Fig. 3a, 3b and 3c).

Talin can adopt multiple conformations in the cell, ranging from a head-to-tail auto-inhibited conformation to a fully extended, mechanically stretched integrin- and actin-bound conformation (Yao et al., 2016). To evaluate the 3D FA distribution of talin as well as its vertical extension, we imaged both N-terminally (talin-1-N) and C-terminally (talin-1-C) tagged talin-1 (Liu et al., 2015; Bouissou et al., 2017). Overall, the z-position values for talin-1-N (Z_centre_=72.8±12) and for talin-1-C (Z_centre_=103.5±20.5nm) are higher than those reported for U2OS cells (Liu et al., 2015). However, the distance between the two ends of talin were comparable to that previously measured and has been considered to represent the extended talin conformation (Case et al., 2015; Liu et al., 2015; Bouissou et al., 2017). Calculations of the inclination angle based on the vertical separation of the two ends and the full-length of the protein, indicated an approximated value of *θ* ≈ 17 °which is also in agreement with previous work (Liu et al., 2015). Thus, our results indicate that talin is fully extended throughout the adhesion and that, in cornerstone FA, talin clusters at higher density at the FA edges (Fig. 2c).

### Vinculin displays a head-above-tail orientation and a cuplike shape distribution

In classical FA, vinculin displays a wide vertical distribution and has a preferred orientation with the tail above the head (Case et al., 2015). iPALM analyses of N-terminally (vinculin-N) and C-terminally (vinculin-C) tagged vinculin constructs (Fig. 3c and Supplementary Fig. 6a) revealed that vinculin vertical positioning in hPSC cornerstone adhesions is significantly higher than in classical FA (Kanchanawong et al., 2010; Case et al., 2015). Increased vinculin z positioning has been linked to vinculin activation and FA maturation (Case et al., 2015), suggesting that hPSC cornerstone adhesions are mature and contain active vinculin. Very unexpectedly, vinculin was oriented “head above the tail” in hPSC cornerstone adhesions (vinculin-N, Z_centre_=100.4±14.4nm; vinculin-C, Z_centre_=76.4±16.3nm) (Fig. 3c). To verify this unexpected observation, we set-up a two-colour iPALM strategy, where Eos-tagged vinculin-C was imaged together with endogenous paxillin (detected with Alexa-647 labelled antibody) at high resolution in the same cells. Using this approach we were able to validate that the vinculin tail is indeed in close relationship with the integrin signalling layer (Fig. 3d). This very peculiar orientation of vinculin was also recently reported in mouse embryonic stem cells FA (Xia et al., 2018, preprint). A “head-above-tail” vinculin conformation could be indicative of an extreme activation state, potentially triggered by the high forces exerted in cornerstone FA (Närvä et al., 2017). Indeed, vinculin has been reported to dissociate from talin in cases where forces borne by an individual protein exceed 25 pN (Yao et al., 2014). In such cases, it is possible that the vinculin head will bind a different partner that localises higher up in cornerstone FA (such as α-actinin-1) or that the vinculin tail engages components close to the plasma membrane such as paxillin or PI(4,5)P_2_ (Carisey and Ballestrem, 2011).

In addition to an inverted orientation, 3D rendering of the acquired localisations for vinculin revealed that its distribution is not homogeneously flat within cornerstone FA but rather that its distribution forms a solid paraboloid or “cup-like” shape (Fig. 3e). This cup-like shape appeared to be specific for vinculin, as it was not observed for paxillin (Fig. 3e), and indicates that membrane distal vinculin molecules occupy a larger lateral area than the membrane proximal vinculin (Fig. 3f). The spatially controlled intra-FA z-distribution of vinculin (Fig. 3e), as well as the ring-shaped distribution of talin and β5 integrin (Fig. 2), suggest that the edge of the pluripotency-associated FA is architecturally unique from its centre.

The unexpected 3D orientation of vinculin in hPSC cornerstone FA prompted us to investigate the role of vinculin in hPSC colony morphology and pluripotency maintenance. We used lentiviral shRNA to silence 95% of endogenous vinculin (Supplementary Fig. 6b). Imaging of control and shvinculin hPSC colonies on VTN showed that, surprisingly, loss of vinculin does not disrupt colony formation, the actin fence or cornerstone FA (Supplementary Fig. 6c and 6d). Vinculin silencing appeared to only modestly increase the size of the small FA found at the centre of hPSC colonies. In addition, silencing of vinculin did not affect Oct4 expression levels (Supplementary Fig. 6b) indicating that vinculin is not required for the maintenance of pluripotency under these conditions.

### Cornerstone adhesions have two vertically separated actin layers

Cornerstone FA are connected by prominent ventral actin bundles (Fig. 1 and Supplementary Fig. 1), and therefore actin is found both inside and outside FA. To accurately determine the vertical positioning of actin in hPSC cornerstone adhesions, endogenous paxillin was used as a reference marker and two-colour iPALM was performed (Fig. 4a and 4b). Unlike the proteins in the integrin signalling and the force transduction layers, actin distribution within cornerstone FA was not Gaussian but could instead be described as the sum of two Gaussian distributions (Fig. 4c). This indicates that, much to our surprise, and in contrast to conventional FA nanoscale architecture (Kanchanawong et al.,2010), actin forms two coexisting vertical layers separated, bya ∼ 50 nm gap on top of the cornerstone FA (Fig. 4b and 4c).

This axial actin localization pattern was further confirmed by the distribution of α-actinin-1, a protein typically overlapping with actin (Carisey et al., 2013). α-actinin-1 also presented the same two-peak distribution than actin albeit with a slight shift in z positioning (Fig. 4d and 4e). Importantly, the separation between the two actin peaks and the two α-actinin-1 peaks was identical suggesting that each actin layer has a corresponding α-actinin-1 layer. The vertical position of the first actin (Z_centre_=99.6±14.4nm) and α-actinin-1 (Z_centre_=125.5±15.1nm) peaks matches the z position reported for these proteins in U2OS cells (Kanchanawong et al., 2010). In contrast, the second peak for actin (Z_centre_=155.4±14.2nm) and α-actinin-1 (Z_centre_=174.8±17.1nm) represents a higher-localising actin population (Fig. 4f), not described before. In the future, it would be interesting to determine the mechanisms regulating this vertical separation of the actin cytoskeleton, and which proteins are positioned in between the two distinct actin layers, within the unusual ∼50 nm gap.

### Nanoscale localization of focal adhesion scaffold proteins kank1 and kank2

While pluripotency FA display 3D stratification of integrin signalling, force transduction and actin regulatory layers, albeit with substantial deviations from those described previously (Liu et al., 2015; Case et al., 2015; Kanchanawong et al., 2010), there is a clear distinction in protein distribution profiles between the FA edge and centre. Therefore, we proceeded to search for possible scaffold proteins that could uniquely mediate the segregation of the cornerstone FA. Two recent reports have described that evolutionary conserved scaffold proteins kank1 and kank2 localise to the border of FA and form links with talin and the actin cytoskeleton (Sun et al., 2016; Bouchet et al., 2016).

We performed two-colour iPALM of Eos-tagged kank1 and kank2 together with endogenous paxillin to enable precise determination of kank 3D localization with respect to FA (Fig. 5a-d). Kank1 localised predominantly at the border of paxillin-positive FA with individual molecules detected inside adhesions (Fig. 5a, 5b). This localisation was validated with endogenous staining and high-resolution Airyscan imaging of kank1 (Supplementary Fig. 3). In contrast, kank2 was always present at the outer-rim and/or outside paxillin-positive FA in hPSC (Fig. 5c). In addition, kank2 only accumulated near cornerstone FA, always towards their distal end, and occasionally following the direction of ventral stress fibres (Fig. 5c). Vertically, both kank1 and kank2 localisation varied depending on their proximity to FA. At the outer-rim of FA, both kank1 and kank2 displayed a higher z position than paxillin (60 nm above paxillin), corresponding to the height of the force transduction layer (Video 2, Fig. 5a, 5c, 5d). Interestingly, in paxillin-negative structures, both kank1 and kank2 localised at a lower vertical position indicating a close proximity to the plasma membrane (Fig. 5b-5d). The z positions obtained for kank1 and kank2 adjacent to FA were (Z_centre_=116.1±12.1nm) and (Z_centre_=100.8±13.4nm) and for FA distal kank1 and kank2 (Z_centre_=63.4±13.6nm) and (Z_centre_=74.2±17.7nm) respectively (Fig. 5e).

**Fig. 5.**
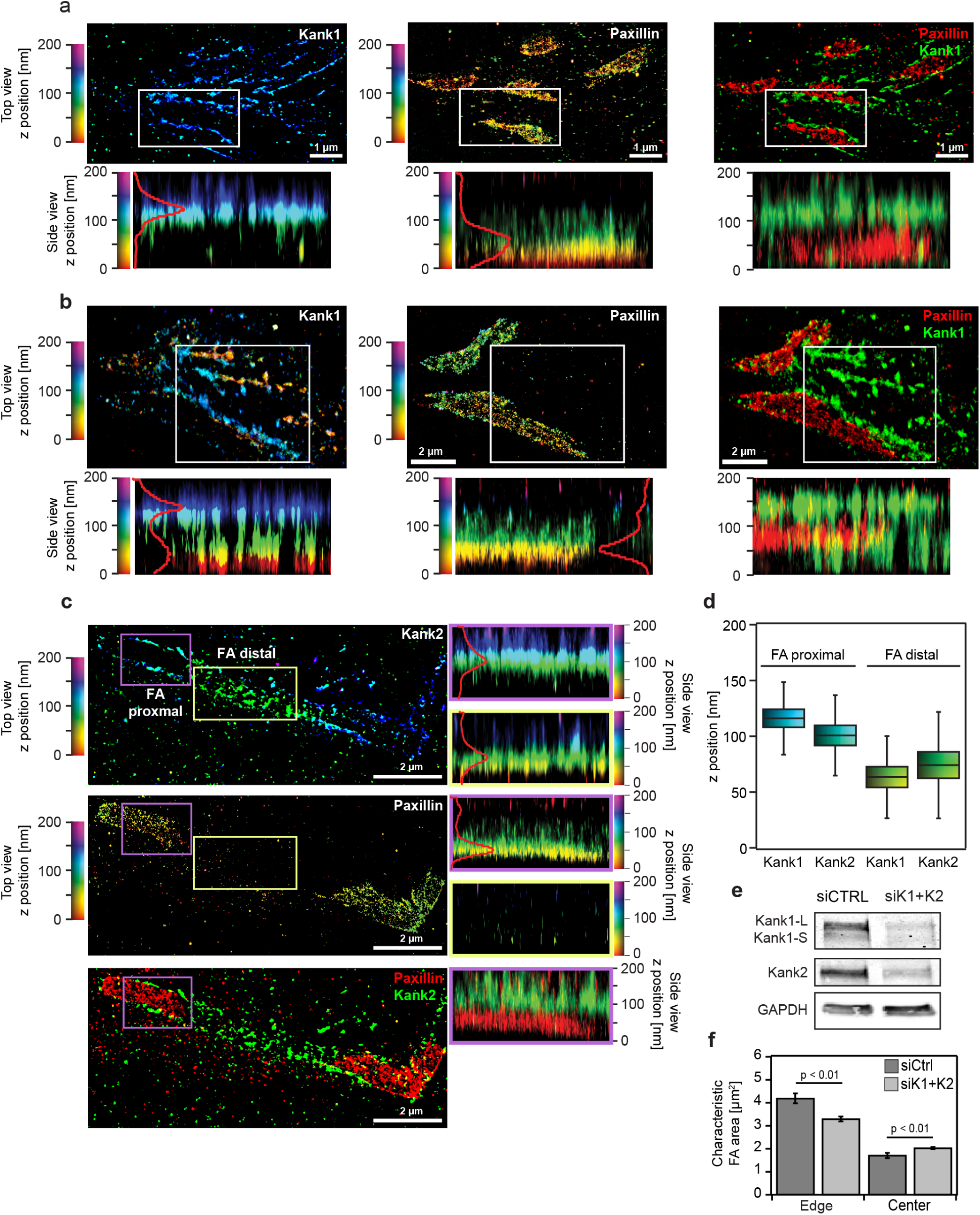
Nanoscale 3D localization of kank1 and kank2 in hPSC. (a-c) Two-colour iPALM images of endogenous paxillin together with Eos-tagged kank1 (a-b) or kank2 (c) in the vicinity of hPSC cornerstone adhesions. Kank1 forms a wall surrounding cornerstone adhesions (a) and is also found outside of adhesions (b). Kank2 also localises both in proximity (c, purple box), and distal (c, yellow box), to cornerstone adhesions. Where localisation of kank1 and kank2 are displayed separately, top-view and side-view images are colour-coded as a function of the z-position of the indicated molecules. Where fluorescence channels are merged (paxillin, red; kank1/kank2, green), the z position is only displayed in the side view and the colours represent the fluorescence signal for each protein. Side views are for selected regions (white, yellow and purple insets). Scale bars 1 μm. (d) iPALM analysis of the Z_centre_ of kank1 and kank2 when in close proximity (adjacent) to cornerstone adhesions or when distant to cornerstone adhesions. (e) Western blot analysis of kank1 and kank2 proteins levels in hPSC pretreated with either control siRNA (siCTRL) or a combination of siRNA targeting kank1 and kank2 (siK1 + siK2) (n=3). (f) Quantification of the characteristic FA area at the edges or in the middle (centre) of hPSC colonies pretreated with siCTRL or siK1 + siK2 (n = 3).

Kank proteins modulate adhesion size and properties in somatic cells by both regulating integrin function and microtubule targeting (Rafiq et al., 2018, preprint). As both kank1 and kank2 localise at the outer-rim of hPSC cornerstone adhesions, we wanted to investigate the contribution of kank1 and kank2 to cornerstone adhesion properties. Unexpectedly, downregulation of kank1 and kank2 expression using siRNA led to smaller cornerstone FA at the colony edge, and, conversely, to larger FA at the centre of the hPSC colonies (Fig. 5e, 5f and Supplementary Fig. 7). Thus, kank1 and kank2 are important regulators for intra-colony distinction of colony centre and edge cells and very interestingly differentially regulate the dynamics of the large cornerstone FA and the smaller adhesions in the colony centre.

## Discussion

An important prerequisite for maintaining hPSC pluripotency in vitro is cell adhesion to specific ECM molecules such as VTN (Braam et al., 2008), highlighting the fundamental role of the adhesion machinery in promoting stemness. We recently described that hPSC colonies are encircled by a strong contractile actin fence connected by exceptionally large FA termed cornerstone FA (Narva et al., 2017). These adhesions are aligned to the colony edge and exert high traction forces upon the substrate (Narva et al., 2017). Here, we investigated the role of these cornerstone adhesions in regulating pluripotency and characterised their 3D architecture using superresolution microscopy.

Our detailed analyses of cornerstone FA, using multiple microscopy approaches, revealed the unique properties of these adhesions. Using live-cell imaging of endogenously-tagged paxillin we demonstrate remarkably slower dynamics, and thus higher stability, of cornerstone FA compared to FA formed at the centre of hPSC colonies. In addition, we demonstrate here that the size and orientation of cornerstone FA are essential for pluripotency, since disrupting the ability of cells to control these parameters using nanopatterns is sufficient to alter colony morphology and trigger differentiation (Fig. 1). These results identify cornerstone FA as guardians of pluripotency in vitro and indicate that their overall architecture is key to their function.

Superresolution iPALM imaging of core FA proteins revealed well-organized vertically demarcated, functional layers within hPSC cornerstone FA (Fig. 6a and 6b). Congruent with classical FA (Kanchanawong et al., 2010), these layers include a membrane-proximal integrin signalling layer composed of integrins and paxillin and a force transduction layer composed of talin and vinculin. The components of the integrin signalling layer (integrins β5 and αV, and paxillin) have a z position indicating a close relationship with the cell membrane and their vertical distribution values are very similar to those reported for U2OS FA (Fig. 6a) (Kanchanawong et al., 2010; Liu et al., 2015; Case et al., 2015). However, the z position of talin-1 and vinculin, components of the force transduction layer, were found to be higher than previously reported for U2OS FA (Kanchanawong et al., 2010) (Fig. 6a). Unlike classical FA, cornerstone FA feature an unprecedented bi-phasic actin regulatory layer (defined by actin and α-actinin-1), where two discrete vertical actin layers are separated by a less actin dense 50 nm gap. This gap may constitute a functional layer in its own right and future work will aim at identifying which proteins localise there. In addition, it will be interesting to determine how these two actin layers are regulated and structurally organised.

**Fig. 6.**
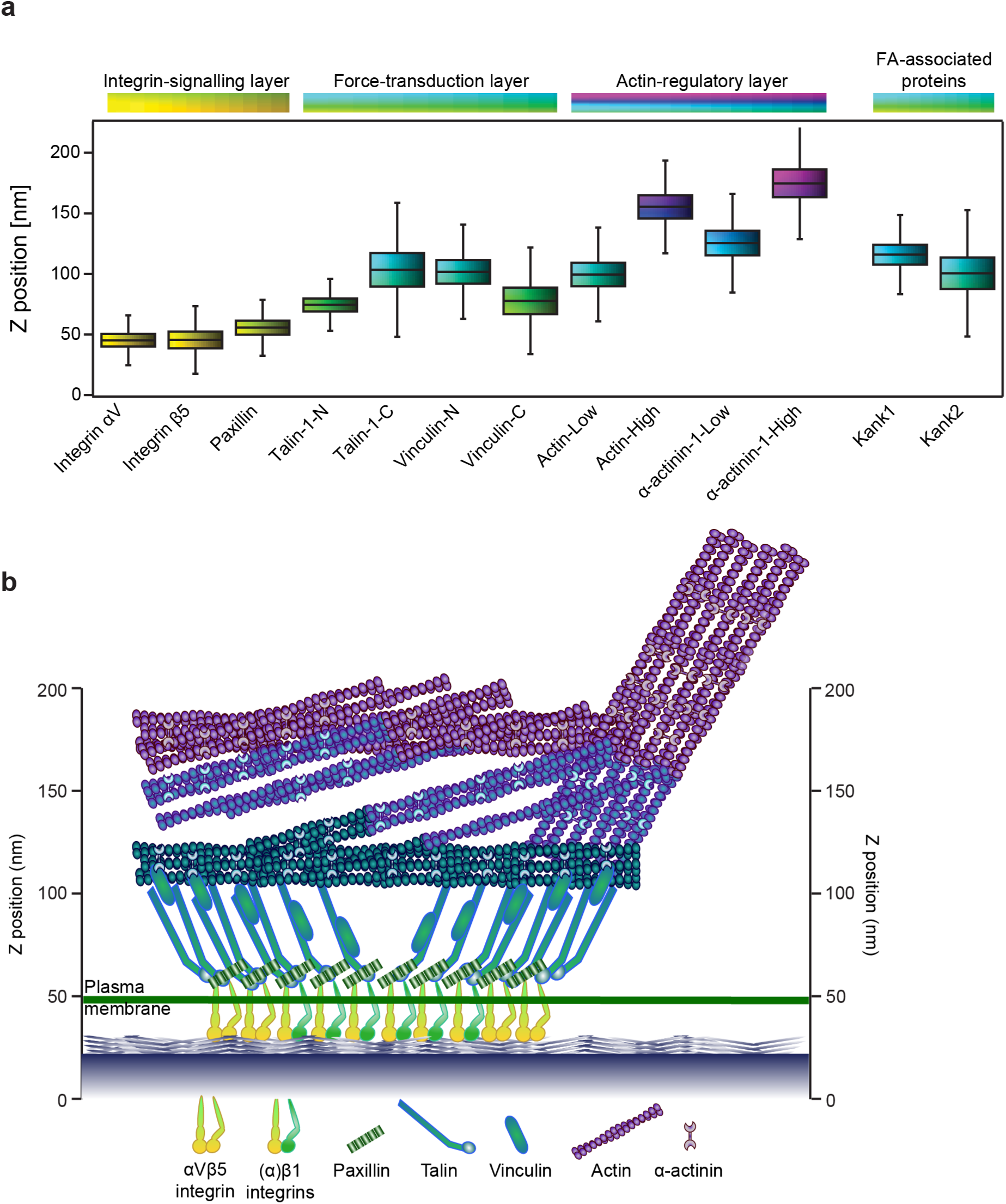
Overall vertical stratification of cornerstone FA. (a) Z positioning (Z_centre_) of αV and β5 integrins, paxillin, vinculin, talin-1, actin, α-actinin-1, kank1 and kank2 in cornerstone adhesions. Letters -N and -C denote the location of the tag while “low” and “high” denote separate peaks in the distribution of the same protein. Results are displayed as boxplots as in figure 2f. (d) Schematic model of the 3D architecture of hPSC cornerstone adhesions. The lateral and vertical positioning of each protein are based on the data presented here. This model does not depict protein stoichiometry.

Here, we also determined for the first time the 3D organization of kank1 and kank2 with respect to the core adhesion. As previously reported for classical FA (Sun et al., 2016; Bouchet et al., 2016), both kank1 and kank2 were laterally distributed around cornerstone FA. However, 3D localisations of kank1 and kank2 revealed that they assemble strong FA-edge-defining vertical “walls” at the outer-rim of paxillin-positive FA rather than a belt (Sun et al., 2016; Bouchet et al., 2016). The vertical positioning of kanks around FA indicate that they are in close proximity to the force transduction layer, which is consistent with the ability of kanks to bind to talin (Bouchet et al., 2016). However, further away from the adhesions both kank1 and kank2 localised vertically much lower, suggesting that recruitment of kanks to FA edges is distinct to their assembly to extended membrane proximal protein assemblies outside of FA. Functionally, kanks regulate the colony centre small FA distinctly from the colony edge cornerstone FA. Kank silencing induced larger and more prominent FA in the colony centre, which is in line with previous work suggesting that in fibroblasts kank2 functions to limit integrin activity and facilitate adhesion sliding (Sun et al., 2016). It is therefore extremely intriguing that the loss of kanks triggers the opposite outcome at the colony edge, reducing cornerstone FA size. Perhaps kanks represent a mechanism that allows hPSC colonies to distinguish between colony edge and centre cells. This concept and the regulatory pathways recruiting kanks to the different hPSC adhesions remain to be investigated.

Detailed imaging also revealed a lateral segregation of specific proteins within the cornerstone adhesions. In particular, β5 integrin was found to assemble high-density ring-like clusters at the edges of adhesions while β1 and αV integrins were found to be more homogeneously distributed. Within classical FA, distinct integrin heterodimers display different dynamics (Rossier et al., 2012) or cluster differently depending on their activation state (Spiess et al., 2018). However, the spatial segregation of specific integrin heterodimers appears to be a unique feature of cornerstone adhesions. This interesting centre-edge segregation within cornerstone FA was also observed with other FA components. Talin density was highest at the FA edges and the vertical distribution of vinculin was higher at FA edges compared to the centre. This apparent intra-FA organisation could be linked to the exceptionally large size of the cornerstone FA and specific interactions of FA components with proteins localising to the outer rim of the FA (FA belt) such as kanks (Bouchet et al., 2016; Sun et al., 2016). Altogether these data indicate that the edges of cornerstone adhesions are structurally distinct from the centre and it is tempting to speculate that the edge and the centre may serve different biological functions, acting either as separate entities or working in synergy to maintain pluripotency of hPSC colonies.

## Materials and Methods

### Reagents, antibodies and compounds

Mouse primary antibodies used in this study were against paxillin (BD biosciences, 612405, 1:1000 for western blotting (WB), 1:100 for immunofluorescence (IF)), β-actin (Sigma Aldrich, Clone AC-15, Cat. No. A1978), GAPDH (Hytest, 5G4MaB6C5, 1:5000 for WB), Sox2 (R&D systems, MAB2018, 1:1000 for WB), vinculin (Sigma Aldrich, V9131, 1:1000 for WB, 1:100 for IF), talin (Sigma Aldrich, clone 8D4, T3287, 1:1000 for WB, 1:100 for IF), β 1 integrin (clone 12G10, in-house production) and α-actinin (Sigma Aldrich, A5044, 1:100 for IF). Rabbit primary antibodies used in this study were against Oct3/4 (Santa Cruz, sc-9081, 1:1000 for WB, 1:100 for IF), kank1 (Bethyl, A301-882A, 1:1000 for WB, 1:100 for IF), kank2 (Sigma aldrich, HPA015643, 1:1000 for WB), integrin β5 (Cell Signaling Technology, 3629, 1:100 for IF) and paxillin (Santa Cruz, sc-5574, 1:100 for IF and for iPALM). The goat anti-Nanog antibody was from R&D systems (AF1997, 1:100 for IF). The secondary antibodies used in this study were all from Thermo Fischer Scientific and used at 1:300. These antibodies included Alexa Fluor 488 donkey anti-rabbit IgG, Alexa Fluor 488 donkey anti-mouse IgG, Alexa Fluor 568 donkey anti-rabbit IgG, Alexa Fluor 568 donkey anti-mouse IgG, Alexa Fluor 647 donkey anti-goat IgG and Alexa Fluor 647 goat anti-mouse IgG. SiR-actin was provided by Cytoskeleton (Cat. No. CY-SC001). Atto 405 phalloidin and Alexa Fluor 488 phalloidin were provided by Thermo Fisher Scientific.

### Plasmids

The following plasmids were provided by Addgene (gift from, Addgene plasmid number): tdEos-Talin-18 (Michael Davidson, 57672) (Kanchanawong et al., 2010), tdEos-Talin-N-22 (Michael Davidson, 57673) (Kanchanawong et al., 2010), tdEos-Vinculin-14 (Michael Davidson, 57691) (Kanchanawong et al., 2010), mEos2-Vinculin-N-21 (Michael Davidson, 57439), tdEos-Lifeact-7 (Michael Davidson, 54527) (Kanchanawong et al., 2010), mEos2-Actin-7 (Michael Davidson, 57339) (Kanchanawong et al., 2010), mEos2-Alpha-Actinin-19 (Michael Davidson, 57346) (Kanchanawong et al., 2010), tdEos-Paxillin-22 (Michael Davidson, 57653), pSpCas9(BB)-2A-GFP (Feng Zhang, 48138) (Ran et al., 2013), AICSDP-1:PXN-EGFP (The Allen Institute for Cell Science, 87420) (Roberts et al., 2017).

The plasmid encoding β5-integrin-mEOS2 was a kind gift from Martin Humphries (University of Manchester, UK). GFP-kank1 and GFP-kank2 plasmids were gifts from Reinhard Fässler (Max Planck Institute of Biochemistry, Martinsried, DE) (Sun et al., 2016).

### Cell culture, gene silencing and transient transfection

The human induced pluripotent stem cell (hPSC) line HEL24.3 was obtained from the University of Helsinki. Cell lines were created using Sendai viruses (Mikkola et al., 2013; Trokovic et al., 2015). Cells were grown in feeder-free conditions on Matrigel (MG) (Corning, 354277) or 5 μg/ml vitronectin (VTN) (Life Technologies, A14700) in Essential 8 (E8) Basal medium (Life Technologies, A15169-01) supplemented with E8 supplement (Life Technologies, A1517-01) at +37°C, 5% CO_2_ in a humidified incubator. Culture media was changed daily. For passaging, cells were detached using sterile-filtered 0.5 mM EDTA (Life Technologies, 15575-038) in PBS for 3 min at room temperature (RT) (Närvä et al., 2017). The parental fibroblast cell line of HEL24.3 was obtained from the University of Helsinki, and were cultured in Dulbecco’s Modified Eagle’s Medium: Nutrient Mixture F-12 (DMEM/F-12) (Life Technologies, 10565-018) supplemented with 10% fetal bovine serum (FCS) (Biowest, S1860) at +37°C, 5% CO2 in a humidified incubator. Fibroblasts were passaged using 0.25% trypsin-EDTA in HBSS (Biowest, L0931-100). U2OS osteosarcoma cells here were grown on VTN coated plates in E8 media. U2OS cells were purchased from DSMZ (Leibniz Institute DSMZ-German Collection of Microorganisms and Cell Cultures, Braunschweig DE, ACC 785).

To induce stem cell differentiation, hPSC were cultured on VTN-coated plates and subsequently differentiated using a mixture of Essential 6 (E6) (Gibco, A1516401) media and 100 ng/ml BMP-4 (R&D systems, 314-BP). The differentiation media was changed daily and the samples were collected after 72 h.

Plasmids of interest were transfected using DNA-in reagent (MTI-Global Stem) according to the manufacturer instructions. Briefly, 24 h before transfection, the cells were seeded on MG to reach appropriate cell confluence of 80%. DNA-in reagent and plasmid DNA were mixed in a ratio of 1:3 (μg of DNA : βl of regent), in Opti-MEM reduced serum medium (Gibco, 31985070), 15 min before the transfection. Cells were gently detached using 0.5 mM EDTA in PBS, diluted in E8 and mixed with transfection mixture with gentle pipetting. The cell-transfection mixture was then added to VTN-coated glass and incubated for 20-24 h.

To transiently suppress kank1 and kank2 expression in hiPSC, Accell siRNA pools (Dharmacon, kank1 E-012879-00-0005; kank2 E-027345-00-0005) were used according to the manufacturer’s protocol. siRNA were diluted to a final concentration of 1 μM. siRNA transfection was done first in suspension during the cell passage and repeated after 24h on adherent cells. 48h after the initial transfection, cells were plated on VTN-coated plates and grown for 24h before fixation or lysis to collect protein samples. siRNA used as a control was Accell Non-targeting Pool (Dharmacon, D-001910-10-20).

To stably suppress vinculin expression, hiPSC were transduced with shRNA containing viral particles in suspension during the cell passage. Puromycin selection (2 μg/ml) was started 48 h after the transduction. The shRNA used as control (shCTRL) was shScramble provided by Sigma Aldrich (SHC002, sequence: CCG GCA ACA AGA TGA AGA GCA CCA ACT CGA GTT GGT GCT CTT CAT CTT GTT GTT TTT). The shRNAs targeting vinculin shVin#1 (TRCN0000116755, sequence : CCG GCG GTT GGT ACT GCT AAT AAA TCT CGA GAT TTA TTA GCA GTA CCA ACC GTT TTT G) and shVin#2 (TRCN0000116756, sequence: CCG GGC TCG AGA TTA TCT AAT TGA TCT CGA GAT CAA TTA GAT AAT CTC GAG CTT TTT G) were provided by the functional genomics unit of the University of Helsinki.

### Generation of C-terminally tagged kank1 and kank2 tdEos constructs

XhoI/EcoRI restriction sites were introduced to flank both mouse kank1 and kank2 genes using polymerase chain reaction (PCR) (Phusion hot start II polymerase, Thermo Fisher Scientific). Primers used were kank1_For (5’-AAT ACT CGA GAT GGC TTA TAC CAC AAA AGT TAA TG-3’) kank1_Rev (5’-AAT AGA ATT CCG TCA AAA GAA CCT CGG TGA G-3’), kank2_For (5’-AAT ACT CGA GAT GGC CCA GGT CCT GC-3’) and kank2_Rev (5’-AAT AGA ATT CCC TCC TCG GCT GAA GAC GA-3’). Following PCR amplification, kank1 and kank2 sequences were inserted into tdEos backbone using XhoI/EcoRI restriction sites. The constructs were sequenced to validate integrity.

### Generation of a paxillin-GFP cell line

The endogenously tagged paxillin HEL24.3 hiPSC line was generated as previously described (Roberts et al., 2017). Briefly, the gRNA sequence targeting paxillin (5’-GCACCTAGCAGAAGAGCTTG-3’) was introduced into pSpCas9(BB)-2A-GFP backbone using the BbsI restriction site. The template plasmid (AICSDP-1:PXN-EGFP) was provided by Addgene (Roberts et al., 2017). HEL 24.3 hPSC were then transfected with the GFP-Cas9-paxillin gRNA construct and template plasmid (AICSDP-1:PXN-EGFP) in equimolar ratio (1:1). After transfection, the cell line was cultured for 5 days before sorting the cells based on green fluorescence with FACS (FACSAria IIu, BD).

### Nanopattern experiments

hPSC were seeded on 35mm nanopattern surface topography glass dishes (Nanosurface Biomedical) coated with 5 μg/ml VTN. The nanopattern topography consisted of several parallel 800 nm wide ridges interspersed with 600 nm deep grooves. Cells were cultured on nanopatterns for 24 h for microscopy in E8 medium or for 3 days for protein analysis in E6 medium. Samples were then acquired either fixing the cells with 4% PFA in PBS for microscopy or lysing the cells for protein analysis.

### SDS-PAGE and quantitative western blotting

Protein extracts were separated under denaturing conditions by SDS-PAGE and transferred to nitrocellulose membranes. Membranes were blocked for 1 h at RT with blocking buffer (LI-COR Biosciences) and then incubated overnight at 4 °C with the appropriate primary antibody diluted in blocking buffer. Membranes were washed with PBS and then incubated with the appropriate fluorophore-conjugated secondary antibody diluted 1:5,000 in blocking buffer for 2 h. Membranes were washed in the dark and then scanned using an Odyssey infrared imaging system (LI-COR Biosciences). Band intensity was determined by digital densitometric analysis using Odyssey software.

### Immunofluorescence

Cells were seeded on VTN-coated glass-bottom microscopy dishes for 20-24 h, washed twice with PBS and simultaneously fixed and permeabilized using 4% PFA and 0.3% Triton-X-100 in PBS for 15 min. Cells were then washed with PBS and the fixative quenched with 0.1 M Glycine for 10 min. following quenching, cells were washed and incubated with primary antibodies diluted in 1% BSA in PBS for either 2 h at RT or at +4°C overnight. The samples were then washed twice with PBS, incubated with secondary antibodies diluted in 1% BSA in PBS for 2 h at RT, washed with 0.2% Tween 20 for 10 min at RT followed by a final PBS wash.

#### Microscopy

The confocal microscope used was a laser scanning confocal microscope LSM880 (Zeiss) equipped with an Airyscan detector (Carl Zeiss). Objectives used were 40x water (NA 1.2) and 63x oil (NA 1.4). The microscope was controlled using Zen Black (2.3) and the Airyscan was used in standard super-resolution mode.

The spinning disk microscope used was a Marianas spinning disk imaging system with a Yokogawa CSU-W1 scanning unit on an inverted Zeiss Axio Observer Z1 microscope controlled by SlideBook 6 (Intelligent Imaging Innovations, Inc.). Objectives used were a 20x (NA 0.8 air, Plan Apochromat, DIC) objective (Zeiss), a 63x oil (NA 1.4 oil, Plan-Apochromat, M27 with DIC III Prism) objective (Zeiss), or a 100x (NA 1.4 oil, Plan-Apochromat, M27) objective. Images were acquired using an Orca Flash 4 sCMOS camera (chip size 2,048 x 2,048; Hamamatsu Photonics).

The structured illumination microscope (SIM) used was DeltaVision OMX v4 (GE Healthcare Life Sciences) fitted with a 60x Plan-Apochromat objective lens, 1.42 NA (immersion oil RI of 1.514), used in SIM illumination mode (five phases x three rotations). Emitted light was collected on a front illuminated pco.edge sCMOS (pixel size 6.5 μm, readout speed 95 MHz; PCO AG) controlled by SoftWorx.

The total internal reflection fluorescence (TIRF) microscope used was a Zeiss Laser-TIRF 3 Imaging System equipped with a 63x (1.46 oil objective, *α* Plan-Apochromat, DIC) objective. Images were acquired on an EMCCD camera (ImageEM C9100-13; chip size 512 x512; Hamamatsu Photonics) controlled by Zen software (Zen 2012 Blue Edition Systems; Zeiss).

### iPALM

iPALM imaging was performed as previously described (Kanchanawong et al., 2010; Shtengel et al., 2009). Briefly, cells were cultured on 25 mm diameter round coverslips containing gold nanorod fiducial markers (Nanopartz, Inc.), passivated with a ca. 50 nm layer of SiO_2_, deposited using a Denton Explorer vacuum evaporator. After fixation, an 18 mm coverslip was adhered to the top of the sample and placed in the iPALM. mEos tagged samples were excited using 561 nm laser excitation at ca. 1-2 kW/cm^2^ intensity in TIRF conditions. Photoconversion of mEos was performed using 405 nm laser illumination at 2-10 W/cm^2^ intensity. 40,000-80,000 images were acquired at 50 ms exposure, and processed/localized using the PeakSelector software (Janelia Research Campus). Alexa Fluor 647 labeled samples were imaged similarly, but with 2-3 kW/cm^2^ intensity 647 nm laser excitation, and 30-40 ms exposure time in STORM-buffer containing TRIS-buffered glucose, glucose oxidase, catalase, and mercaptoethanol amine (Dempsey et al., 2011).

iPALM data were analysed and images rendered using the PeakSelector software (Janelia Research Campus) as previously described (Kanchanawong et al., 2010; Shtengel et al., 2009). iPALM localization data records both the fluorescent molecules localized within the focal adhesions (FA) as well as molecules in the cytoplasmic fraction, to quantify the spatial distribution of the proteins within individual FA, we zoomed into areas covered only by the FA and the immediate surrounding space. To render iPALM images, a single colour scheme was used from red to purple, covering the z range 0-200 nm, where features within FA are seen. The same colour scheme was also used for side-view (xz) images. For analysis of protein distributions in FA: Z_centre_ and svert calculation, the three-dimensional molecular coordinates for each region (individual FA) were analysed to obtain histograms of vertical positions with 1-nm bins. The centre vertical positions (Z_centre_) and width (svert) were determined from a Gaussian fit to the FA molecule peak. For proteins like actin and α-actinin where dual peaks were observed, the fitting was done using the sum of two-Gaussian distributions with independent centre vertical position and width. After the histograms for all images and individual FA were obtained, they were combined into a single average Z_centre_ and svert.

### Quantification of focal adhesion properties

For the quantification of FA area, FA number, density and characteristic area were analysed using the ImageJ software (NIH) in a similar way as previously described (Närvä et al., 2017). In short, individual FA were first identified and measured using the ‘analyze particles’ built-in function of ImageJ. Second, as we observed that the distribution of FA area was not Gaussian but rather a heavy-tailed distribution, this was approximated, and fitted in Igor Pro (WaveMetrics) to a hyper-exponential distribution or, in other words, a weighted sum of two exponential densities. Finally, the fit provided us with two characteristic areas of FA for each curve, A1 describing the smaller FA and A2 describing the larger FA. As A1 was not significantly different between samples, we focused on the differences found on A2 that describes the larger FA rarely present in fibroblasts or at the center of hPSC colonies, but regularly present at the edge of hPSC colonies. For simplicity, throughout the paper the characteristic area A2 is referred to as ‘characteristic FA area’. In the case of hPSC colonies, as there were clear visible differences in the FA size at the edge of the colony, we separated the edge from the center for quantification. The edge was defined as a 40-pixel-thick strip starting from the line delimiting the colony boundary and extending towards the inside of the colony. The centre was defined as the remaining region of the colony.

## Statistical analysis

If not indicated otherwise, statistical analyses were performed using an analysis of variance (ANOVA) complemented by Tukey’s honest significant difference test (Tukey’s HSD). The software R version 3.3.3 (R Development Core Team, Vienna, Austria) was used to perform these analyses. Statistical significance levels are annotated as ns = nonsignificant (p > 0.05) or by providing the range of the p-value.

## Data availability

The authors declare that the data supporting the findings of this study are available within the article and from the authors on request.

## Acknowledgements

We thank J. Siivonen and P. Laasola for technical assistance. H. Hamidi is acknowledged for editing the manuscript and the figures. The Ivaska lab is acknowledged for lively discussions and critical feedback on the manuscript. We thank Michael Davidson, Feng Zhang, Reinhard Fässler, Janet Askari, Martin Humphries and The Allen Institute for Cell Science for providing reagents. The Cell Imaging Core (Turku Centre for Biotechnology, University of Turku, Åbo Akademi University and Biocenter Finland) and the functional genomics unit of the University of Helsinki (Research Programs Unit, HiLIFE Helsinki Institute of Life Science, Faculty of Medicine, University of Helsinki, Biocenter Finland) are acknowledged for services, instrumentation, and expertise.

iPALM imaging was performed in collaboration with the Advanced Imaging Center at Janelia Research Campus, a facility jointly supported by the Gordon and Betty Moore foundation and the Howard Hughes Medical Institute.

This study has been supported by the Academy of Finland (G.J., E.N. and J.I.), Academy of Finland CoE for Translational Cancer Research (J.I.), ERC CoG grant 615258, Sigrid Juselius Foundation and the Finnish Cancer Organization (J.I.). A.S. has been supported by the University of Turku Doctoral programme for Molecular Medicine (TuDMM). iPALM microscopy was performed at the Howard Hughes Medical Institute (HHMI) Janelia Research Campus Advanced Imaging Center (AIC). The AIC is jointly sponsored by the HHMI and the Gordon and Betty Moore Foundation.

## Conflicts of interests

The authors declare no competing financial interests.

## Authors contributions

Conceptualization, J.I., C.G. and A.S.; Methodology, A. S., C.G., E.N., J.A., G.J. ; Formal Analysis, C.G., A.S., E.N., J.A.; Investigation, A.S., C.G., E.N., J.A., M.S., J.I.; Resources, M.M.; Writing-Original Draft, C.G., J.I., G.J.; Writing - Review and Editing, C.G., A.S., E.N., J.I., G.J.; Visualization, C.G., A.S., G.J.; Supervision, J.I. and T-L.C.; Funding Acquisition, J.I.

